# RNAPosers: Machine Learning Classifiers For RNA-Ligand Poses

**DOI:** 10.1101/702449

**Authors:** Sahil Chhabra, Jingru Xie, Aaron T. Frank

## Abstract

Determining the 3-dimensional (3D) structures of ribonucleic acid (RNA)-small molecule complexes is critical to understanding molecular recognition in RNA. Computer docking can, in principle, be used to predict the 3D structure of RNA-small molecule complexes. Unfortunately, retrospective analysis has shown that the scoring functions that are typically used to rank poses tend to misclassify non-native poses as native, and *vice versa*. This misclassification of non-native poses severely limits the utility of computer docking in the context pose prediction, as well as in virtual screening. Here, we use machine learning to train a set of pose classifiers that estimate the relative “nativeness” of a set of RNA-ligand poses. At the heart of our approach is the use of a pose “fingerprint” that is a composite of a set of atomic fingerprints, which individually encode the local “RNA environment” around ligand atoms. We found that by ranking poses based on the classification scores from our machine learning classifiers, we were able to recover native-like poses better than when we ranked poses based on their docking scores. With a leave-one-out training and testing approach, we found that one of our classifiers could recover poses that were within 2.5 Å of the native poses in ∼80% of the 88 cases we examined, and similarly, on a separate validation set, we could recover such poses in ∼70% of the cases. Our set of classifiers, which we refer to as RNAPosers, should find utility as a tool to aid in RNA-ligand pose prediction and so we make RNAPosers open to the academic community via https://github.com/atfrank/RNAPosers.

## Introduction

Beyond acting as an intermediary between deoxyribonucleic acid (DNA) and proteins, ribonucleic acids (RNAs) play key regulatory roles within the cell^1–3^. For instance: ribosomal RNAs (rRNAs) catalyze protein synthesis^4^; riboswitches turn on and off RNA transcription or translation^5^; and short interfering RNAs (siRNAs)^6^ and microRNAs (miRNAs)^7^ silence the expression of targeted mRNAs. Indeed, many classes of “functional” RNAs are implicated in diseases^8^ and are now considered viable drug targets^9–12^. Moreover, targeting RNAs with small molecules has garnered keen interest over the last decade^13–16^.

Rational structure-based methods promise to be a viable approach for identifying small molecules that can bind to and modulate the activity of therapeutically relevant RNAs. Crucial to the success of rational structure-based approaches in RNA drug discovery is the ability to accurately predict the 3-dimensional (3D) structure of the complex formed between an RNA and a small molecule ligand. In principle, computer docking algorithms can be used to predict the 3D orientation and conformation (referred to as the pose) of a ligand bound to an RNA receptor. Unfortunately, “redocking” tests reveal that state-of-the-art scoring functions typically fail to recover the correct poses^17–21^. In this respect, there is an urgent need for methods that can accurately distinguish “native-like” RNA-ligand poses from non-native decoy poses.

Recently, machine learning has been used to address several challenges associated with computer docking and virtual screening. For protein-ligand complexes in particular, machine learning has been used to develop more robust scoring functions for both pose and binding affinity prediction.^22–26^ Here, we used machine learning to train a set of pose classifiers that quantify the “nativeness” of RNA-ligand complexes. In what follows, we summarize our comparison between the ability of docking scores and machine learning classifiers to rank and identify atomically correct RNA-ligand poses. Compared with docking scores, we found that machine learning pose-classifiers were better able to discriminate native-like RNA-ligand poses from decoy poses.

## MATERIALS AND METHODS

### Decoy sets

We compiled an initial dataset comprised of 88 RNA-ligand systems. An additional set of 17 RNA-ligand system was compiled and used for final validation. For both datasets, the crystal structures of the RNA-ligand complexes were downloaded from the Protein Data Bank (PDB:http://www.pdb.org). To generate diverse decoy sets for each RNA-ligand system, computer docking was performed using the docking program rDock^27^. The following protocol was used to generate the poses with rDock (Figure 1A and B). First, a set of poses were generated in the actual binding pocket, using the reference ligand method, with the sphere radii from the center of the known binding pocket set to 2, 3, 4, 5, 6, 7, and 8 Å, respectively. At each sphere radius, 50 poses were generated, for a total of ∼350 poses. Next, 250 additional poses were generated by docking into the binding pockets that were identified using the two-sphere method, with outer sphere radii set to 20, 40, 60, 80 and 100 Å, respectively. Hence, in total, ∼600 poses were generated for each RNA-ligand complex. For some RNA-ligand complexes, the number of poses were less than 600 because the two-sphere method failed to identify binding pockets at one or more of the outer sphere radii we utilized for binding pocket detection. All pocket detection was carried out using the rDock utility program, rbcavity. The entire set of decoy poses can be accessed at https://github.com/atfrank/RNAPosers.

**Figure 1.**
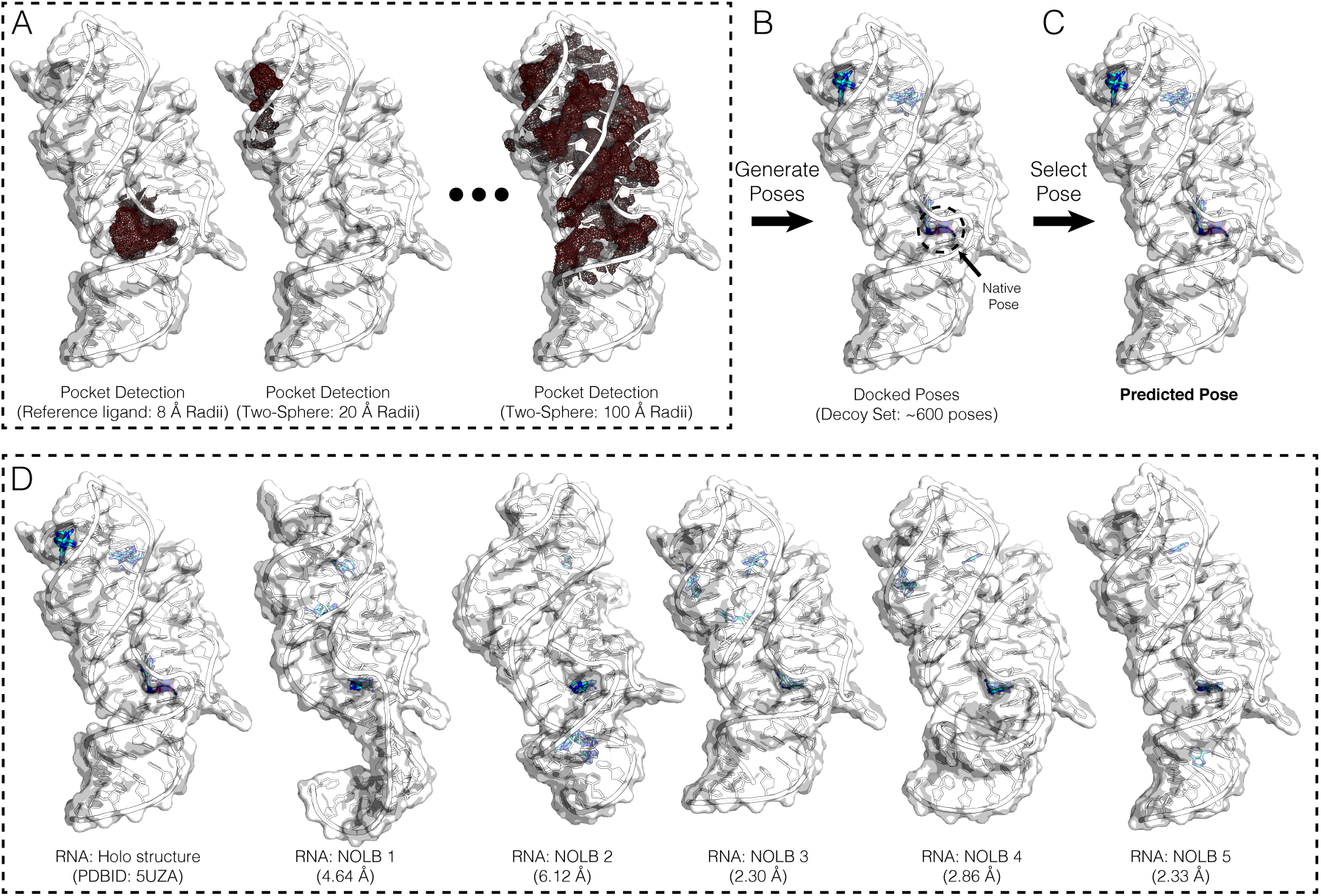
Illustrated are the steps involved in generating the decoy sets used in this study. (A) First, the actual binding pocket is mapped using the reference ligand method, and second, alternative pockets are mapped using the two-sphere methods, with increasingly large radii. (B) Third, poses were generated by docking the ligands into each of the mapped binding pockets and then combining all poses into a single decoy set. (C) The focus of this study was to develop and assess methods for selecting atomically-correct poses from these decoy poses. (D) Example of an augmented decoy set in which both the RNA and ligand are flexible. For a given RNA-ligand complex, we generated the augmented decoy sets by deforming the holo RNA structure along its non-linear normal modes. In this study, we generated five such structures for each RNA and then generated docked poses for each, and then all poses combined to form a final augmented decoy set. Shown here is that augmented decoy set for PDBID: 5UZA. Indicated under each of the deformed structures is the RMSD relative to the holo structure.

### Pose classifiers

Machine learning was used to train a set of pose classifiers that take a set of “pose features” as input and output a measure of the “nativeness” of the pose. First, we generated a set of classifiers for which the “pose features” correspond to individual scoring terms in the rDock scoring function.^27^ Second, we generated a set of classifiers for which the “pose features” correspond a pose novel fingerprint the depends on the pairwise distance between heavy atoms in the an RNA receptor and a the heavy atoms in a small molecule ligand (see below). To train the pose classifiers, we employed the random forest method implemented in the sklearn Python module.^28^ The classifiers comprised of an ensemble of 1000 decision trees with class weight set to balanced subsample. All other parameters were set to their default values. The classifiers were trained using a leave-one-out approach using the set of poses generated using rDock (see above). We trained separate classifiers with the nativeness RMSD thresholds set to ≤1.0, 1.5, 2.0, and 2.5Å. Machine learning models can be susceptible to the so-called “twinning effect,” which occurs when samples in the training set closely resemble samples in testing set. Here we have employed leave-one-out cross-validation in an attempt to mitigate the potential impact of “twinning” when assessing the performance of classifiers. In this leave-one-out approach, a single RNA-ligand system was removed from the training set and the classifiers were trained on the remaining 87 RNA-ligand complex. The resulting classifier was then assessed on the excluded RNA-ligand system. *If the ligand in any of the other 87 RNA-ligand systems was identical to the ligand in the left-out system, they were removed prior to training the classifier used to assess the left-out system.*

### Pose fingerprint

We utilized a pose fingerprint that is a composite of a set of atomic fingerprints. For a given ligand atom, the atomic fingerprint correspond to the vector, {*V*_*i*_}, whose elements *V*_*i*_(*η, v*) are given by

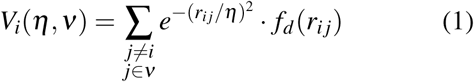

where *r*_*i j*_ is the distance between the heavy atom *i* in a ligand and the heavy atom *j* in the RNA receptor, *η* is the width of a Gaussian function (here we set *η* = 2), *ν* is a set of unique RNA atom types, and *f*_*d*_(*r*_*ij*_) is the damping function given by

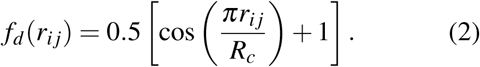

Here, *R*_*c*_ is a cutoff distance and in this study, it was set to 20 Å. We note that the atomic fingerprint based on Eq. 1, which is a multi-element extension of the atomic fingerprint developed by Botu and et.al.^29^, is invariant to the basic atomic transformation operations of translation, rotation and permutation.

For a given ligand pose, *i*, a fingerprint vector, *F*_*i*_, was generated from the atomic fingerprint defined by Eq. 1 by summing over all instances of a given atompair type, which is defined by the SYBYL atom types in the ligand and atom types in the RNA. We denote each unique ligand-RNA pair as *S*. As such, an element in the fingerprint for pose *i* and atom pair type *S, F*_*i,S*_, is given by

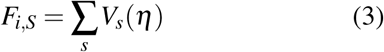

Here *s* runs over all instances of pair type *S* in pose *i*. If pair-type *S* is not present in *i, F*_*i,S*_ = 0. The set of 21 SYBYL atom types we used were: {C.1, C.2, C.3, C.ar, C.cat, N.1, N.2, N.3, N.4, N.ar, N.am, N.pl3, O.2, O.3, O.co2, S.2, S.3, S.o, S.o2, P.3}. The set of 85 RNA atom types we used were: ADE:{C1′, C2, C2′, C3′, C4, C4′, C5, C5′, C6, C8, N1, N3, N6, N7, N9, O2′, O3′, O4′, O5′, OP1, OP2, P}; CYT:{C1′, C2, C2′, C3′, C4, C4′, C5, C5′, C6, N1, N3, N4, O2, O2′, O3′, O4′, O5′, OP1, OP2, P}; GUA:{C1′, C2, C2′, C3′, C4, C4′, C5, C5′, C6, C8, N1, N2, N3, N7, N9, O2′, O3′, O4′, O5′, O6, OP1, OP2, P}; URA:{C1′, C2, C2′, C3′, C4, C4′, C5, C5′, C6, N1, N3, O2, O2′, O3′, O4, O4′, O5′, OP1, OP2, P}. Thus, the final pose fingerprint *F*_*i*_ = {*F*_*i,S*_}, which was normalized for each RNA-ligand system, contained 1785 elements (21 SBYL types × 85 RNA atom types). Coincidentally, our pose fingerprint closely resembles a recently described fingerprint that was successfully used to train machine learning pose and binding affinity predictors.^26^

### Assessing classifiers

In order to quantify our ability to recover atomically correct poses using either docking scores from the rDock scoring function or the classification scores from our pose classifiers, we first sorted the poses. When using docking scores, the pose with *lowest* (most negative) score was then identified and the RMSD relative to crystal pose was determined. When using classification scores, the pose with *highest* classification score was identified and the RMSD relative to crystal pose was determined. We also calculated the success rates *S*(*X*) as the percentage of RNA-ligand complexes for which the RMSD of the best pose (identified using either docking scores or classification scores) were within *X* Å of the corresponding crystal pose.

## Results and Discussion

For protein-ligand complexes, modern scoring functions have a reported success rate that exceeds ∼75 %.^30^ In contrast, for RNA-ligand complexes, state-of-the-art scoring functions have a success rate near 50 %.^21;27^ This discrepancy between the success rate of protein and RNA scoring functions motivated us to explore methods capable of enhancing our ability to discriminate native-like poses from non-native decoys.

### Docking scores exhibit low success rates

We began our study by assessing the ability of docking scores to recover the correct pose from decoy poses located in the experimental binding pocket as well as decoy poses located in alternate pockets on the surface of the RNA. To accomplish this, we initially generated decoys sets comprised of ∼600 diverse poses for 88 RNA-ligand complexes (see Methods; Figure 1A-C). In these decoys sets, the RNA receptors corresponded to the holo structures where only the ligand orientation and conformation varied.

Shown in Figure 2A are the distributions of the RMSD (relative to the crystal pose) of the best poses selected from these decoy sets using individual score terms in the rDock scoring function.^27^ When using the total docking score, the median RMSD of the predicted pose was 3.41 Å (Figure 2A; Table 1). We obtained similar results when using the total interaction, the van der Waals interaction, and the polar interaction score terms. In these cases, the median RMSD were 5.72, 4.75, and 6.88 Å, respectively. To better quantify the ability of the score terms to select atomically correct poses, we also computed the success rate, *S*(*X*), defined as the percentage of cases in which the predicted pose was within *X* Å of the native pose. Using the total docking score, the *S*(1.00), *S*(1.50), *S*(2.00), and *S*(2.5) were 22.7, 29.5, 37.5, 42.0, and 44.3 %, respectively (Table 1). Similarly, the *S*(1.00), *S*(1.50), *S*(2.00), and *S*(2.5) were 17.0, 21.6, 27.3, and 33.0%, respectively, when using total interaction, 18.2, 22.7, 28.4 and 36.4%, respectively, when using the van der Waals interaction, and 8.0, 9.1, 12.5, and 21.6%, respectively, when using the polar interaction score terms (Table 1). The docking score terms in the rDock scoring function, therefore, exhibited marginal ability to recover correct poses from diverse decoys poses.

**Table 1.**
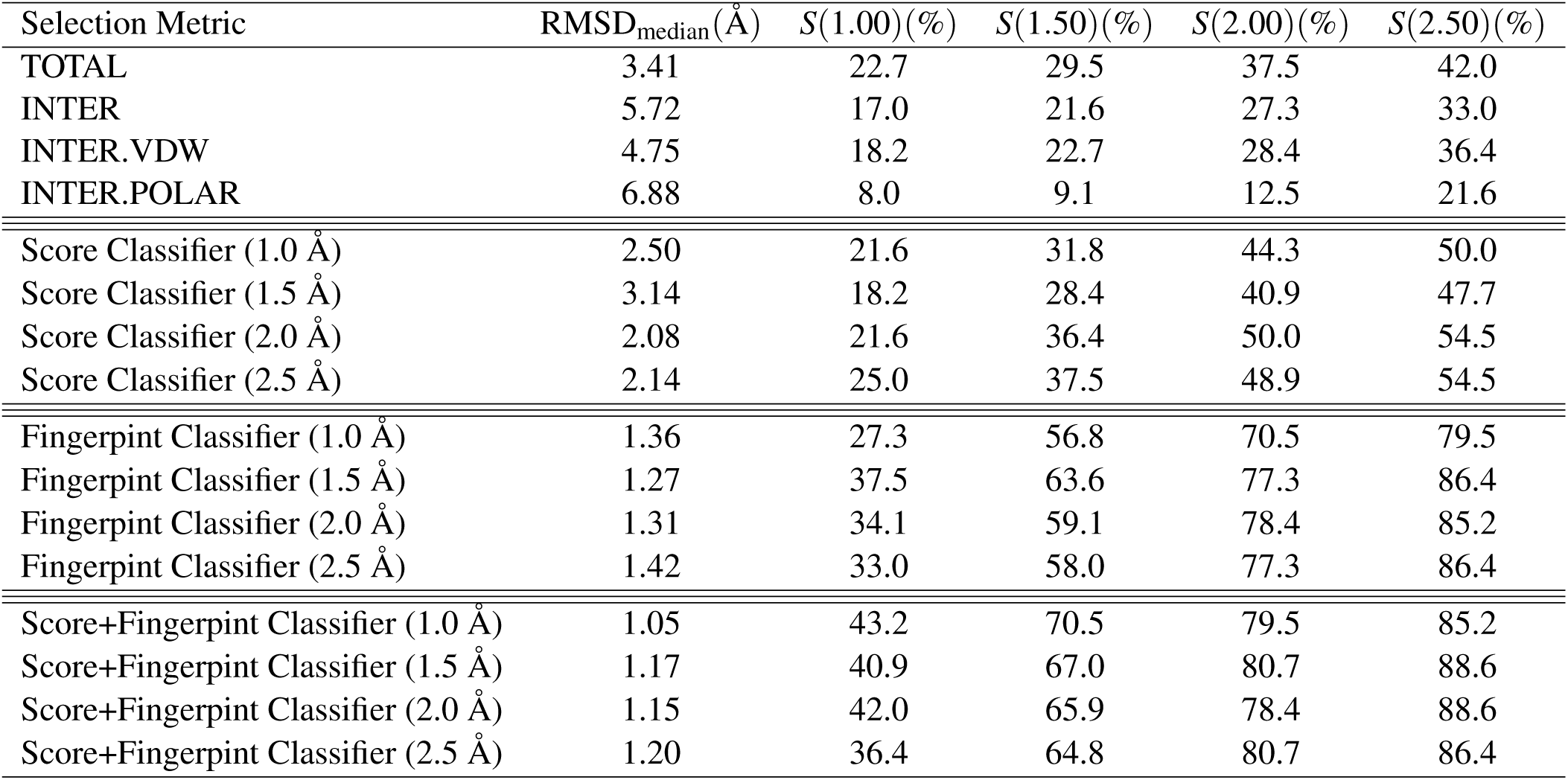
Median RMSD and success rates for systems in the leave-one-out training database. Listed are the results obtained when the best poses were selected using the docking score terms and classifiers that were trained using the docking score terms, our pose fingerprint, and docking scores plus our pose fingerprint as learning features. For the pose classifiers, we include results for classifiers that we trained with the nativeness threshold set to 1.0, 1.5, 2.0, and 2.5 Å.

| Selection Metric | RMSD<sub>median</sub>(Å) | $S(1.00)$(%) | $S(1.50)$(%) | $S(2.00)$(%) | $S(2.50)$(%) |
| --- | --- | --- | --- | --- | --- |
| TOTAL | 3.41 | 22.7 | 29.5 | 37.5 | 42.0 |
| INTER | 5.72 | 17.0 | 21.6 | 27.3 | 33.0 |
| INTER.VDW | 4.75 | 18.2 | 22.7 | 28.4 | 36.4 |
| INTER.POLAR | 6.88 | 8.0 | 9.1 | 12.5 | 21.6 |
| Score Classifier (1.0 Å) | 2.50 | 21.6 | 31.8 | 44.3 | 50.0 |
| Score Classifier (1.5 Å) | 3.14 | 18.2 | 28.4 | 40.9 | 47.7 |
| Score Classifier (2.0 Å) | 2.08 | 21.6 | 36.4 | 50.0 | 54.5 |
| Score Classifier (2.5 Å) | 2.14 | 25.0 | 37.5 | 48.9 | 54.5 |
| Fingerprint Classifier (1.0 Å) | 1.36 | 27.3 | 56.8 | 70.5 | 79.5 |
| Fingerprint Classifier (1.5 Å) | 1.27 | 37.5 | 63.6 | 77.3 | 86.4 |
| Fingerprint Classifier (2.0 Å) | 1.31 | 34.1 | 59.1 | 78.4 | 85.2 |
| Fingerprint Classifier (2.5 Å) | 1.42 | 33.0 | 58.0 | 77.3 | 86.4 |
| Score+Fingerprint Classifier (1.0 Å) | 1.05 | 43.2 | 70.5 | 79.5 | 85.2 |
| Score+Fingerprint Classifier (1.5 Å) | 1.17 | 40.9 | 67.0 | 80.7 | 88.6 |
| Score+Fingerprint Classifier (2.0 Å) | 1.15 | 42.0 | 65.9 | 78.4 | 88.6 |
| Score+Fingerprint Classifier (2.5 Å) | 1.20 | 36.4 | 64.8 | 80.7 | 86.4 |

**Figure 2.**
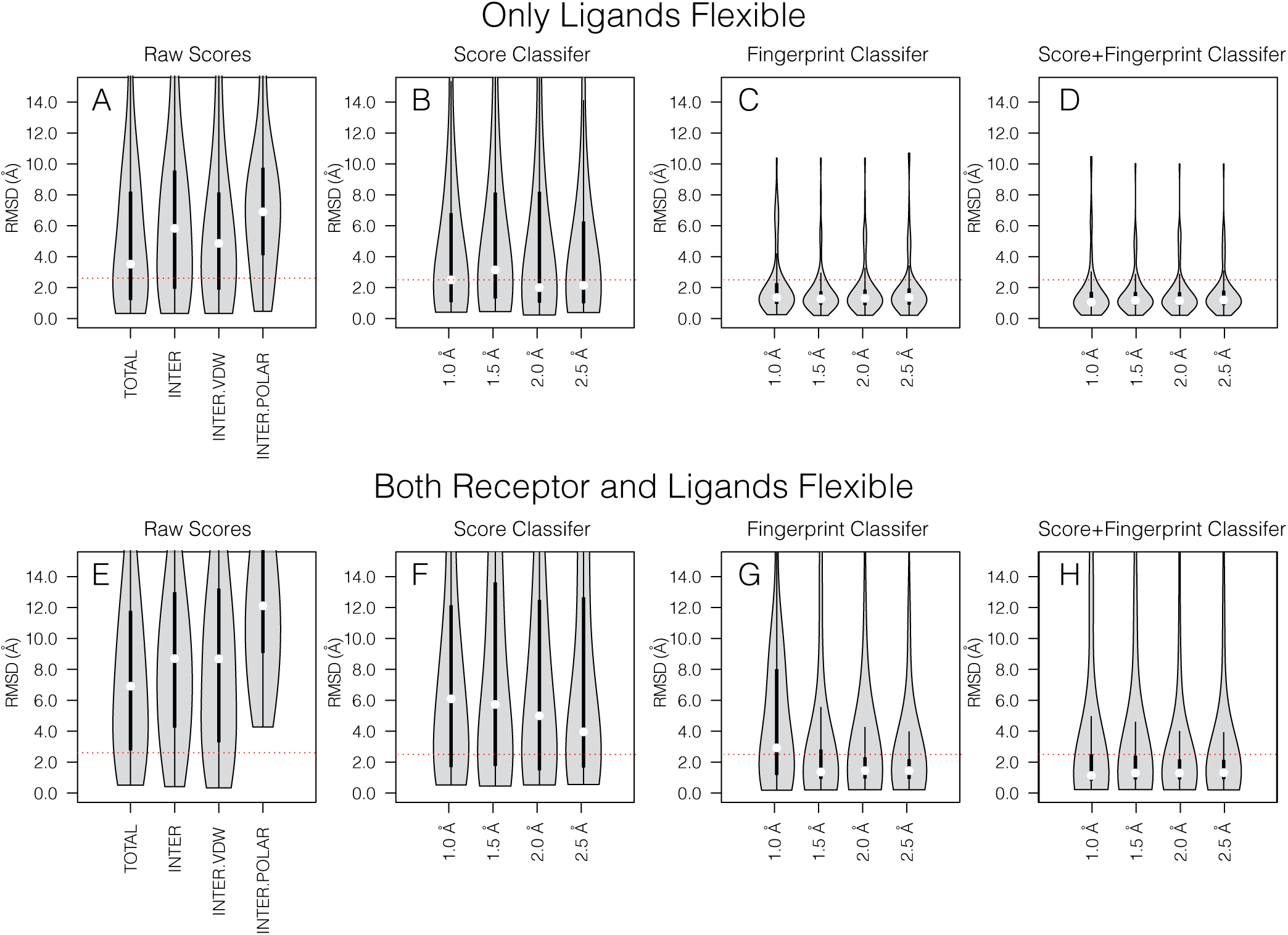
RMSD distributions of the best predicted poses over the systems in the leave-one-out training database when the best poses were predicted using (A) docking score terms and classifiers training using the (B) docking score terms, (C) our pose fingerprint, and (D) raw docking scores and our pose fingerprint as features, respectively. For the pose classifiers, results are shown for independent sets of classifiers that were trained with the nativeness threshold set to 1.0, 1.5, 2.0, and 2.5 Å.

### Pose classifiers improve success rates

Next, we asked whether nonlinear classifiers could enhance our ability to recover the correct poses from decoy poses. To test this, we cast the problem of recovering correct ligand poses as a “classification” problem and then machine learning models were trained to discriminate correct poses from decoy poses. Briefly, we built a set of random forest classification models that take a set of features as input and output “classification scores” that estimate the probability of a pose being native-like. To accomplish this, we first trained a series of random forest pose classifiers using a leave-one-out cross-validation approach in which we selected a single RNA-ligand from the dataset of 88 RNA-ligand systems (the leave-one-out dataset), and trained a classifier using the decoy sets for the remaining 87 RNA-ligand systems. After training, the performance of the resulting classifier was assessed using the RNA-ligand system set of the left-out system. For that system, the classification scores for all decoy poses were determined and then the pose with the highest classification score was selected as the “best” (or predicted) pose for the left-out system. This procedure was repeated 88 times, i.e., one for each system in the leave-one-out dataset.

Shown in Figure 2B are the distributions of the RMSD of the predicted poses that were identified using the classifiers that used the individual terms in the rDock scoring function as learning features. Reported are results for the classifiers trained with nativeness RMSD threshold set to 1.0, 1.5, 2.0, and 2.5 Å, respectively. Listed in Table 1 are the corresponding success rates. In general, the RMSD of the best poses that were identified using the score-based pose classifiers were lower than those of the best poses selected using the terms in the rDock scoring function. For instance, for the score-based pose classifiers trained with nativeness threshold set to 1.0, 1.5, 2.0, and 2.5 Å, the the median RMSD of the best poses were 2.50, 3.14, 2.08, and 2.14, respectively (Figure 2B; Table 1). The success rates, *S*(1.00), *S*(1.50), *S*(2.00), and *S*(2.5) were also generally larger for the score-based classifiers, with the best results obtained with the nativeness threshold set to 2.0 and 2.5 Å, respectively. *S*(1.00), *S*(1.50), *S*(2.00), and *S*(2.5) were 21.6, 36.4, 50.0, and 54.5%, respectively, for the classifiers trained with the threshold set to 2.0 Åand 25.0, 37.5, 48.9, and 54.5%, respectively, for the classifiers trained with the threshold set to 2.5 Å. In comparison, the values obtained when using the total docking score to identify the best pose were 37.5, 42.0, and 44.3 %, respectively (Table 1). These results suggest that the pose classifiers that were trained using the scores terms as learning features could boost our ability to recover correct poses. The success rates, however, still pales in comparison to the success rates of protein-ligand pose prediction methods, several of which achieve success rates near 75%.

As such, we next asked whether we could further enhance the success rate of RNA-ligand pose prediction by training pose classifiers on features that more directly depend on RNA-ligand interactions. Specifically, we were interested in examining the utility of a simple distance-based atomic fingerprint that describes the local atomic environment near a given site which has shown promise in predicting properties like atomic forces^31^ and resembles a pose fingerprint recently used for protein-ligand pose predictions.^26^ To create a composite fingerprint from atomic fingerprints, we summed and normalized all atomic fingerprints associated with specific ligand-RNA pair “types” (see Methods and Fig. 1). Using this composite RNA-ligand interaction fingerprint, we then trained another set of pose classifiers, again using the leave-out-one cross-validation approach. For comparison, we also trained classifiers that used the rDock score terms plus our pose fingerprint as features. Here again, separate classifiers were trained with the nativeness threshold set to 1.0, 1.5, 2.0, and 2.5 Å.

For the pose fingerprint classifiers trained with nativeness threshold set to 1.0, 1.5, 2.0, and 2.5 Å, the median RMSD of the best poses were 1.36, 1.27, 1.31, and 1.42 Å, respectively (Figure 2C; Table 1). These fingerprint-based classifiers all exhibit similar success rates. For instance, *S*(1.00), *S*(1.50), *S*(2.00), and *S*(2.5) were 37.5, 63.6, 77.3, and 86.4 %, respectively, for the classifiers trained with the nativeness threshold set to 1.5 Å, and which had the lowest median RMSD of 1.27 Å(Table 1). In comparison, *S*(1.00), *S*(1.50), *S*(2.00), and *S*(2.5) were 33.0, 58.0, 77.3, and 86.4 %, respectively, for the classifiers trained with the nativeness threshold set to 2.5 Å, and which had the highest median RMSD of 1.42 Å(Table 1). We obtained comparable results for the pose classifiers that were trained using the docking scores plus the fingerprint as features. Notable among these was the classifier trained with the nativeness threshold set to 1.0 Å; for this set of classifiers, the median RMSD of the best poses was 1.05 Åand the *S*(1.00), *S*(1.50), *S*(2.00), and *S*(2.5) were 43.2, 70.5, 79.5, and 85.2, respectively. Based on this leave-one-out analysis, the pose classifiers trained using the pose fingerprint as well as the classifiers trained using docking score terms and pose fingerprint as features, both exhibited remarkable ability to recover atomically correct poses from the leave-one-out decoy sets.

### Pose classifiers improve success rates on augment decoys sets in which both the RNA and the ligand are flexible

As a more robust test of our classifiers, the original decoy set for each RNA-ligand system in our dataset was augmented with poses that were generated by docking against a set of five perturbed structures of the corresponding RNA receptor. As might be expected, the individual docking score terms failed to recover poses in these augmented decoy sets, with the best results obtained when ranking and selecting poses based on their total docking score. In this case, the median of RMSD of the best pose was 6.81 Å (Figure 2E; Table 2). Marginally better results were obtained using the score classifiers (Figure 2F; Table 2), with the classifiers trained with a nativeness threshold of 2.5 Å exhibiting the lowest median value of 3.96 Å, and *S*(1.00), *S*(1.50), *S*(2.00), and *S*(2.5) values of 13.8, 22.5, 32.5, and 37.5 %, respectively (Table 2). By contrast, the pose classifiers trained using the pose fingerprint as features and the composite score and pose fingerprint features were typically able to recover correct poses (Figure 2G and H). Except for the fingerprint classifier trained with the nativeness threshold set to 1.00 Å, the median RMSD for the predicted poses, were *<*1.50 Å. A representative of these pose classifiers was the fingerprint classifier trained with nativeness threshold set to 2.50 Å. For this classifier, the median RMSD of the predicted poses was 1.44 Å and the success rates, *S*(1.00), *S*(1.50), *S*(2.00), and *S*(2.5), were 31.2, 53.8, 72.5, and 82.5 %, respectively (Table 2).

**Table 2.** Median RMSD and success rates for the augmented decoy sets (Figure 1D) of the systems in the leave-one-out training database. Listed are the results obtained when the best poses were selected using the docking score terms and classifiers that were trained using the docking score terms, our pose fingerprint, and docking scores plus our pose fingerprint as learning features. For the pose classifiers, we include results for classifiers that we trained with the nativeness threshold set to 1.0, 1.5, 2.0, and 2.5 Å.

| Selection Metric | RMSD<sub>median</sub>(Å) | $S(1.00)$(%) | $S(1.50)$(%) | $S(2.00)$(%) | $S(2.50)$(%) |
| --- | --- | --- | --- | --- | --- |
| TOTAL | 6.81 | 6.2 | 11.2 | 18.8 | 23.8 |
| INTER | 8.59 | 2.5 | 6.2 | 11.2 | 16.2 |
| INTER.VDW | 8.58 | 6.2 | 10.0 | 15.0 | 21.2 |
| INTER.POLAR | 12.00 | 0.0 | 0.0 | 0.0 | 0.0 |
| Score Classifier (1.0 Å) | 6.10 | 15.0 | 21.2 | 30.0 | 33.8 |
| Score Classifier (1.5 Å) | 6.02 | 15.0 | 21.2 | 31.2 | 37.5 |
| Score Classifier (2.0 Å) | 5.00 | 15.0 | 25.0 | 37.5 | 41.2 |
| Score Classifier (2.5 Å) | 3.96 | 13.8 | 22.5 | 32.5 | 37.5 |
| Fingerprint Classifier (1.0 Å) | 2.98 | 20.0 | 30.0 | 38.8 | 46.2 |
| Fingerprint Classifier (1.5 Å) | 1.35 | 33.8 | 53.8 | 63.8 | 72.5 |
| Fingerprint Classifier (2.0 Å) | 1.42 | 31.2 | 53.8 | 72.5 | 80.0 |
| Fingerprint Classifier (2.5 Å) | 1.44 | 31.2 | 53.8 | 72.5 | 82.5 |
| Score+Fingerprint Classifier (1.0 Å) | 1.10 | 41.2 | 62.5 | 70.0 | 76.2 |
| Score+Fingerprint Classifier (1.5 Å) | 1.31 | 36.2 | 57.5 | 68.8 | 76.2 |
| Score+Fingerprint Classifier (2.0 Å) | 1.30 | 35.0 | 57.5 | 70.0 | 80.0 |
| Score+Fingerprint Classifier (2.5 Å) | 1.32 | 31.2 | 56.2 | 72.5 | 77.5 |

Next, we repeated the analysis described above for an independent dataset comprised of 17 RNA-ligand systems that we did not include in the leave-one-out dataset used to train the pose classifiers. These tests were carried out using the more challenging augmented decoys sets. When using the docking scores to rank and select poses, the van der Waals interaction scores exhibited the lowest median RMSD poses (3.29 Å; Table 3) and when using the score-based pose classifiers, the classifier trained with nativeness threshold of 1.0 Å exhibited the lowest median RMSD (4.19 Å; Table 3). In contrast, the corresponding values for the fingerprint-based classifiers and scores plus fingerprint-based classifiers were 1.89 Å (nativeness threshold of 1.00 Å) and 2.05 Å (nativeness threshold of 2.50 Å), respectively (Table 3). In terms of the success rates, the best performing classifiers were the fingerprint-based pose classifiers that were trained with nativeness threshold of 1.00 and 1.50 Å, respectively. The success rates, *S*(1.00), *S*(1.50), *S*(2.00), and *S*(2.5) were 35.3, 47.1, 52.9, and 58.8 for the fingerprint-based pose classifier that were trained with nativeness threshold of 1.00 Å and 41.2, 41.2, 47.1, and 64.7 for the corresponding classifier that were trained with nativeness threshold of 1.50 Å (Table 3). As such, though the success rates on the validation set were lower than rates estimated from our leave-one-out analysis, they are significantly higher than the results obtained using either the raw docking scores or the baseline score-based classifiers (Table 2).

**Table 3.** Median RMSD and success rates for the augmented decoy sets (Figure 1D) of the systems in the independent validation set. Listed are the results obtained when the best poses were selected using the docking score terms and classifiers that were trained using the docking score terms, our pose fingerprint, and docking scores plus our pose fingerprint as learning features. For the pose classifiers, we include results for classifiers that we trained with the nativeness threshold set to 1.0, 1.5, 2.0, and 2.5 Å. Also listed are results for a meta classifier that classified pose based on classification scores from the score-based, fingerprint-based, and score and fingerprint-based classifiers, as well as the total docking score and interaction docking score term. For all pose classifiers, results are shown for independent sets of classifiers that were trained with nativeness threshold set to 1.0, 1.5, 2.0, and 2.5 Å.

| Selection Metric | RMSD<sub>median</sub>(Å) | $S(1.00)(\%)$ | $S(1.50)(\%)$ | $S(2.00)(\%)$ | $S(2.50)(\%)$ |
| --- | --- | --- | --- | --- | --- |
| TOTAL | 4.96 | 5.9 | 5.9 | 17.6 | 17.6 |
| INTER | 7.16 | 5.9 | 11.8 | 11.8 | 11.8 |
| INTER.VDW | 3.29 | 5.9 | 17.6 | 17.6 | 23.5 |
| INTER.POLAR | 11.67 | 0.0 | 0.0 | 0.0 | 0.0 |
| Score Classifier (1.0 Å) | 4.19 | 11.8 | 17.6 | 17.6 | 17.6 |
| Score Classifier (1.5 Å) | 6.62 | 5.9 | 17.6 | 17.6 | 17.6 |
| Score Classifier (2.0 Å) | 5.89 | 5.9 | 17.6 | 17.6 | 17.6 |
| Score Classifier (2.5 Å) | 8.39 | 5.9 | 17.6 | 23.5 | 23.5 |
| Fingerprint Classifier (1.0 Å) | 1.89 | 35.3 | 47.1 | 52.9 | 58.8 |
| Fingerprint Classifier (1.5 Å) | 2.05 | 41.2 | 41.2 | 47.1 | 64.7 |
| Fingerprint Classifier (2.0 Å) | 2.69 | 29.4 | 41.2 | 41.2 | 47.1 |
| Fingerprint Classifier (2.5 Å) | 2.12 | 23.5 | 35.3 | 41.2 | 58.8 |
| Score+Fingerprint Classifier (1.0 Å) | 2.69 | 35.3 | 41.2 | 41.2 | 47.1 |
| Score+Fingerprint Classifier (1.5 Å) | 2.71 | 29.4 | 41.2 | 41.2 | 47.1 |
| Score+Fingerprint Classifier (2.0 Å) | 2.69 | 29.4 | 41.2 | 41.2 | 47.1 |
| Score+Fingerprint Classifier (2.5 Å) | 2.05 | 29.4 | 41.2 | 47.1 | 58.8 |
| Meta Classifier (1.00 Å) | 2.53 | 17.6 | 17.6 | 23.5 | 41.2 |
| Meta Classifier (1.50 Å) | 1.56 | 29.4 | 47.1 | 58.8 | 58.8 |
| Meta Classifier (2.00 Å) | 1.68 | 29.4 | 41.2 | 58.8 | 58.8 |
| Meta Classifier (2.50 Å) | 1.68 | 29.4 | 47.1 | 58.8 | 70.6 |

Finally, we examined whether we could further enhance pose prediction by training meta-classifiers that combined the classification scores from the individual score- and fingerprint-based classifiers. This approach commonly referred to as “stacking” or “blending” has been shown to improve the performance of a collection of individual models on common learning task.^32^ To accomplish this, we fitted a simple logistic regression model that estimated the nativeness of a set of poses from the classification scores of those poses that were derived from the set of 12 independent pose classifiers (Table 3) as well as the total docking score, the total van der Waals interaction scores, and the total polar interaction score. We fitted a separate meta-classification models for each of the four nativeness thresholds we used in this study (namely, 1.0, 1.5, 2.0, and 2.5 Å). Each model was trained using the augmented decoy sets (Figure 1D) of the 88 systems used in our initial leave-one-out analysis and tested on the augmented decoy sets of the 17 systems in the validation set. With the exception of the meta-classifier trained with the nativeness threshold set to 1.0 Å, the best poses selected with meta-classifiers all exhibited median RMSD values *<*2.0 Å; the median RMSD were 1.56, 1.68, and 1.68 Å, for the meta-classifiers trained with nativeness threshold set to 1.5, 2.0, and 2.5 Å, respectively, compared to a value of 2.53 for the classifier trained with the nativeness threshold set to 1.0 Å (Table 3). Based on the success rates, the meta-classifier trained with the nativeness threshold set to 2.5 Å exhibited the best results (Figure 3). For that classifier, the *S*(1.00), *S*(1.50), *S*(2.00), and *S*(2.5) were 29.4, 47.1, 58.8, and 70.6, respectively. As such, for the systems in this validation set, we discovered meta pose classifiers enabled us to recovery atomically correct poses from the augmented decoy sets with an accuracy level superior to those of the individual pose classifiers we initially trained and tested.

**Figure 3.**
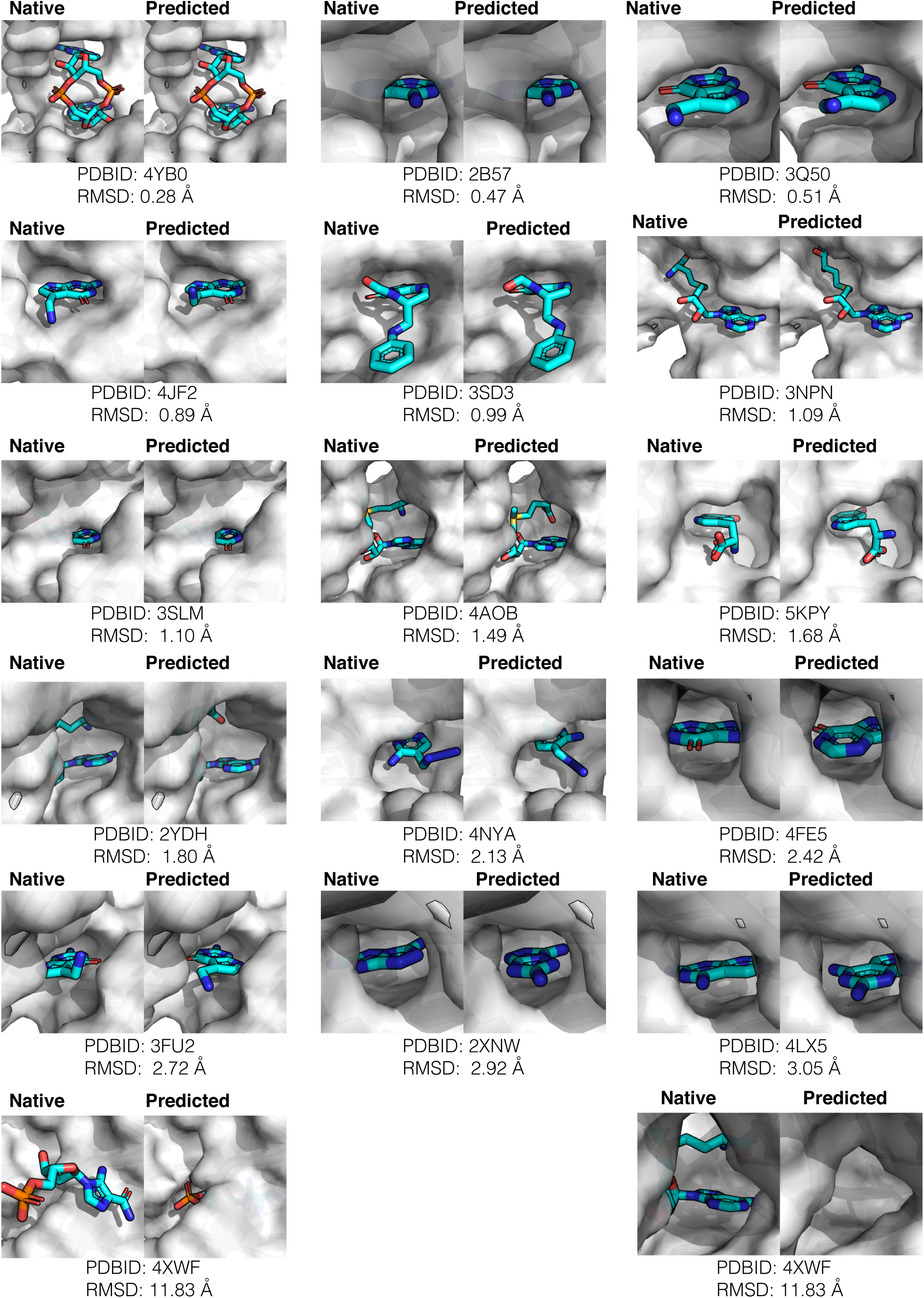
Native versus predicted for systems in the validation set. Shown are visual comparisons between the native poses (*left*) and predicted poses (*right*) that were selected using the best performing meta-classifier, which we trained with nativeness threshold set to 2.5 Å.

When we challenged the classifiers with the augmented decoys sets, in which both the RNA conformation and ligand poses were varied, one possible reason why their overall performance degraded is that we trained the classifiers themselves on decoys sets in which only the ligand poses were varied (i.e., for each RNA-ligand complex, the RNA was fixed in the *holo* conformation). It seems reasonable that the performance of the classifiers could enhanced the by training them using the augmented decoy sets, in which both the RNA conformation and ligand poses were varied. Unfortunately, such augmented decoy sets are *severely* imbalanced, which hampers the training of robust random forest classifiers. In future work, we explore methods to train robust classifiers on these highly imbalanced decoy sets as well as developing similar classifiers to RNA-protein complexes.

Here, we focused on training classifiers that can be used to post-process docked RNA-ligand poses. The enhanced ability to recover atomically accurate poses indicate that the relationships between the pose fingerprint and the nativeness of individual poses captured by our classifiers might also be useful in guiding conformational sampling during docking. In principle, we could convert these classification scores into a pseudo-energy term of the form −*kT* ln(1 + *p*), and can add it as an additional term to existing scoring functions. Alternatively, a new pair-wise scoring function could be developed using the random forest refinement strategy recently described by Merz and coworkers^25^. In either case, we could then assess the classifier-informed scoring functions by quantifying the extent to which the distribution of docked poses shift towards or away from native-like poses. Future work will explore this further.

## Conclusion

In this study, we showed that machine learning classifiers significantly enhance RNA-ligand pose prediction accuracy, especially when applied to set of ligand poses that were docked against the *holo* conformations of RNAs. Due to the promising results we obtained using our pose classifiers, we have incorporated them into the software tool, RNAPosers (https://github.com/atfrank/RNAPosers). To facilitate the development and testing of other RNA-ligand pose prediction methods and scoring functions, we also make accessible all of the decoy sets used in this study. In the context of RNA-ligand pose prediction, RNAPosers should find utility as a tool to assess the relative quality of a set of poses derived either from purely computational methods or from hybrid modeling methods that incorporate experimental data such as chemical shift perturbation. Also, within the context of virtual screening, we envision that RNAPosers may find utility as a tool to identify high-confidence poses that can be brought forward for binding affinity prediction using physics-based free energy calculation methods like, MM-PBSA and FEP calculations.

## Author contributions statement

A.T.F. conceived project, S.C. and J.X. conducted the computational experiments, S.C. and A.T.F. analyzed the results, J.X. implemented the RNAPosers tool, and S.C., J.X., and A.T.F. wrote the manuscript. All authors reviewed the manuscript.

## Acknowledgements

The authors would like to acknowledge the valuable scientific contribution of Sean M. Law, who wrote the C++ library used to implement the pose fingerprint code.

## Notes

https://github.com/atfrank/RNAPosers

